# Two fiber pathways connecting amygdala and prefrontal cortex in humans and monkeys

**DOI:** 10.1101/561811

**Authors:** Davide Folloni, Jérôme Sallet, Alexandre A. Khrapitchev, Nicola R. Sibson, Lennart Verhagen, Rogier B. Mars

## Abstract

The interactions between amygdala and prefrontal cortex are pivotal to many neural processes involved in learning, decision-making, emotion, and social regulation. The broad functional role of amygdala-prefrontal interplay may reflect the diversity of its anatomical connections. Little, however, is known of the structural wiring linking amygdala and prefrontal cortex in humans. Using diffusion imaging techniques, we reconstructed connections between amygdala, anterior temporal and prefrontal cortex in human and macaque brains. First, by studying macaques we were able to assess which aspects of connectivity known from tracer studies could be identified with diffusion imaging. Second, by comparing diffusion imaging results in humans and macaques we were able to estimate amygdala-prefrontal connection patterns in humans and compare them with those in the monkey. We observed a prominent and well-preserved bifurcation of connections between amygdala and frontal lobe into two fiber networks – an amygdalofugal path and an uncinate fascicle path – in both species.

## INTRODUCTION

There has recently been a particular focus on understanding amygdala-prefrontal pathways in health and disease. These neural circuits have been implicated with reward-based learning and decision-making (De Martino et al., 2006; Hunt and Hayden, 2017; Morrison and Salzman, 2010; Murray and Wise, 2010; Rudebeck and Murray, 2014; Rushworth et al., 2011), emotional and social behavior (Noonan et al., 2014; Volman et al., 2011; Whalen et al., 1998). Recent models, therefore, increasingly emphasize how amygdala and prefrontal cortex (PFC) do not encode isolated computations but have instead complementary roles in these processes (Munuera et al., 2018; Saez et al., 2015). To understand these computational roles, however, requires a complementary understanding of the anatomy underlying the brain systems involved (Marr, 1982) which still remains to be established. This necessity is particularly evident in the context of developing new therapeutic approaches for psychiatric disorders. For example, to target effective deep brain stimulation at fine-scale white matter (WM) architecture in the context of obsessive-compulsive disorder or major depressive disorder (Lozano et al., 2019).

Based on WM dissections and tract-tracing studies in non-human species, two pathways are known to connect anterior temporal regions including the amygdala and the frontal lobe: the uncinate fascicle (UF) and ventral amygdalofugal pathway (AmF, from here on also referred to as the ‘amygdalofugal’; Nieuwenhuys et al., 2008; Schmahmann and Pandya, 2006). However, to date, the AmF has not been yet described in great detail in the primate brain (Carmichael and Price, 1995; Croxson et al., 2005; Lehman et al., 2011). In fact, this pathway is often entirely neglected in human studies of circuits for social-emotional behavior and decision making. This is especially striking given its potential to inform directionality of influence: while the UF is a bi-directional tract with many projections from prefrontal cortex terminating in medial and lateral basal nuclei of the amygdala, the AmF predominantly carries efferent projections from the amygdala to subcortical and prefrontal targets (Bickart et al., 2012). Potentially, this dissociation can help to inform whether abnormal neural processes central to certain psychiatric disorders, are more related to efferent or afferent projections of the amygdala (Bramson et al., 2019 biorXiv; Murray and Rudebeck, 2018; Volman et al., 2013). As such, it is critical to examine the degree to which tract-tracing anatomical schemes in non-human species are applicable to the human brain.

Diffusion-weighted magnetic resonance imaging (MRI) and associated tractography algorithms now enable one to make direct inter-species comparison of anatomical networks possible. Strengths of this methodology also include its whole-brain coverage, speed of acquisition, and applicability in many species including humans (Mars et al., 2014). We therefore used diffusion-weighted MRI and tractography to map the pathways connecting amygdalar territory and frontal regions in the macaque monkey and human brains focusing specifically on AmF and UF. Using the same technique in the two species, allows us to directly compare changes in connectional architecture across macaque and human brains.

Apart from a direct comparison of connections between species, macaque diffusion MRI data also served another important purpose. There are limits in the degree to which diffusion MRI tractography can accurately identify origin-to-termination axonal projections, as tract-tracing techniques can (Maier-Hein et al., 2017; Mars et al., 2016a; Thomas et al., 2014). In fact, diffusion MRI tractography is better suited to the complementary goal of reconstructing the body and course of white matter bundles through the cortical white matter (Catani and Thiebaut de Schotten, 2008; Mars et al., 2018). Following this notion, we seed the tractography directly in the white matter core of the fiber bundles of interest, rather than in either the amygdalar or frontal grey matter. Critically, this approach requires detailed anatomical knowledge to guide tractography algorithms; these data are available in the macaque monkey from histology and tracer injection studies.

We therefore used high-resolution diffusion MRI data in the macaque to ascertain the degree to which the diffusion MRI tractography approach identified aspects of amygdala-frontal connectivity known previous macaque studies. This demonstrated that the core bundles of the AmF perforate the porous substantia innominata in the basal forebrain. In this inhomogeneous tissue the fractional anisotropy index is not fully informative of tract integrity. While this could lead to false negatives in deterministic tractography, importantly, the probability distribution of streamlines running through this area is still valid. Accordingly, here we have adopted a probabilistic approach to tractography strongly informed by prior anatomical knowledge to reconstruct this tract in a manner similar to previous tract tracing studies.

Using this approach, we were able to robustly define and dissociate two pathways connecting amygdala and frontal lobe regions in the macaque and human brains: a medial amygdalofugal pathway and an orbital uncinate tract. Together, these pathways form a larger constellation of amygdala-frontal circuits, but each with a distinct connectional profile interfacing with a unique set of brain regions. While the amygdalofugal pathway predominantly connected the amygdala and nucleus accumbens with subgenual cingulate, ventromedial, and frontopolar regions in ventral PFC, the uncinate pathway connected anterior temporal regions, including the amygdala, with predominantly areas in lateral orbital and frontopolar cortex. The relationship and structure of these tracts is preserved across primate species, supporting the translation of insights from non-human primate anatomy to inform our understanding of the human brain.

## METHODS

### Data acquisition

Human in-vivo diffusion MRI data were provided by the Human Connectome Project (HCP), WU-Minn Consortium (Principal Investigators: David Van Essen and Kamil Ugurbil; 1U54MH091657) funded by the 16 NIH Institutes and Centers that support the NIH Blueprint for Neuroscience Research; and by the McDonnell Center for Systems Neuroscience at Washington University (Van Essen et al., 2013). The minimally preprocessed datasets of the first twenty-four subjects from the Q2 public data release were used (age range 22-35 years; thirteen females). Data acquisition protocols are detailed in Ugurbil et al. (2013) and Sotiropoulos et al. (2013). The diffusion data were collected at a 1.25 mm isotropic resolution across the entire brain on a customized 3T Siemens Skyra scanner using a monopolar Stejskal-Tanner diffusion scheme with a slice-accelerated EPI readout. Sampling in q-space included 3 shells at b = 1000, 2000, and 3000 s/mm^2^. For each shell 90 diffusion gradient directions and 6 non-diffusion weighted images (b = 0 s/mm^2^) were acquired with reversed phase-encoding direction for TOPUP distortion correction (Andersson et al., 2003). A subject-specific cortical surface mesh with 10k vertices was created using FreeSurfer on the T1-weighted image (acquired using an MPRAGE sequence at 0.7 mm isotropic resolutions) and aligned to the diffusion space as part of the HCP’s minimum preprocessing pipeline (Glasser et al., 2013).

Macaque diffusion MRI data were obtained *ex-vivo* from three rhesus monkeys (*Macaca mulatta*, age range at death 4-14 years, mean age 8 years, standard deviation 6.6; one female) using a 7T magnet with a Varian DirectDrive^tm^ (Agilent Technologies, Santa Clara, CA, USA). Immediately after death, the brains were perfusion fixed with formalin and stored. Approximately one week before MRI scanning they were perfused in phosphate buffer solution to enhance their diffusion signal. During scanning, the brains were suspended in fomblin™, which has no magnetic properties itself and therefore allows an unbiased scan. For each brain, nine non-diffusion-weighted (b = 0 s/mm^2^) and a single diffusion-weighted (b = 4000 s/mm^2^) datasets were acquired using a single line readout, 2D Stejskal-Tanner pulse sequence (Stejskal and Tanner, 1965) (TE/TR: 25 ms/10 sec; matrix size: 128 x 128; resolution 0.6 mm x 0.6 mm; 128 slices; slice thickness: 0.6 mm; 131 isotropically distributed diffusion directions; receiver bandwidth: 100 kHz). In addition, in the same session for each brain a T1-map, resulting in a T1-weighted image, was collected (MPRAGE sequence at TE/TR = 8ms/10s; Ti = 10.6 ms; Matrix Size = 128 x 128; 128 slices; slice thickness: 0.6 mm). Death and fixation have an impact on the brain, causing a reduction in *e*x*-vivo* tissue diffusivity. Therefore, to achieve an equivalent diffusion contrast to in-vivo data, the diffusion coefficient was increased from b = 1000 to 4000 s/mm^2^ (Mars et al., 2016a) Importantly, although diffusion magnitude is affected and requires compensation, diffusion anisotropy is largely preserved *post-mortem* (D’Arceuil and de Crespigny, 2007).

### Data preprocessing

Human data were preprocessed according to the HCP minimal preprocessing pipelines (Glasser et al., 2013; Sotiropoulos et al., 2013). Macaque data preprocessing and all subsequent analyses were performed using tools from FSL (Smith et al., 2004) and the in-house MR Comparative Anatomy Toolbox (MrCat). In short, non-diffusion-weighted images were extracted from the full set, averaged, and corrected for spatial RF-field inhomogeneity bias before being linearly registered to the average T1-weighted structural image of the same animal. The T1-weighted images in turn were linearly and non-linearly registered to a dedicated ex-vivo macaque T1w template aligned to the standard macaque F99 space (Van Essen and Dierker, 2007). Combined, these registrations allowed a direct mapping between diffusion-, T1-weighted, and standard space for both human and macaque data.

Diffusion tensors were fitted to each voxel using DTIFIT (Behrens et al., 2003). Voxel-wise crossing-fiber model fitting of diffusion orientations was performed in both human and macaque datasets using FSL’s BedpostX (Behrens et al., 2007). For the human in-vivo data, a multi-shell extension was used to reduce overfitting of crossing fibers due to non-monoexponential diffusion decay (Jbabdi et al., 2012). Up to three fiber orientations per voxel were allowed. This produced voxel-wise posterior distributions of fiber orientations that were subsequently used in probabilistic tractography.

### Probabilistic tractography

The principal aim of the tractography analyses was to reconstruct the prefrontal projections of the ventral amygdalofugal pathway (AmF) and uncinate fascicle (UF). Tractography recipes for both were created by reference to previously published atlases (Nieuwenhuys et al., 2008; Paxinos et al., 2000), tract-tracing studies (Fudge et al., 2002; Lehman et al., 2011; Schmahmann and Pandya, 2006), and diffusion tractography studies (Catani et al., 2013; Croxson et al., 2005; Jbabdi et al., 2013). All masks were created in F99 space (Van Essen and Dierker, 2007) in the monkey brain and MNI space in the human brain and subsequently warped to each individual’s diffusion space and adjusted by hand to ensure anatomical accuracy.

The seed mask of the AmF was drawn in the sub-commisural white matter perforating the substantia innominata, a region between the dorsal amygdaloid and the bed nuclei of the stria terminalis (BNST). The mask was aimed to carefully reproduced macaque tract tracing studies (deCampo and Fudge, 2013; Fudge and Haber, 2001; Schmahmann and Pandya, 2006; Mai et al., 2015; Paxinos et al., 2000). In accordance with the local trajectory of the AmF as estimated by tract tracing the seed mask was constrained to contiguous voxels showing high fractional anisotropy in an anterior-posterior direction. UF was seeded axially in the anterior temporal lobe, in the white matter rostro-lateral to the amygdala. After testing various coronal seed masks, an axial seed mask was chosen to account for the strong curve of the fibers from dorsal-ventral to anterior-posterior orientation to orientation as they enter into the frontal lobe (Catani and Thiebaut de Schotten, 2008).

To test the specificity of these tractography approach, we reconstructed three alternative tracts projecting to the prefrontal cortex: the anterior limb of the internal capsule (ICa), the extreme capsule (EmC), and the cingulum bundle (CB). These tracts were chosen because together they constitute prominent systems connecting thalamic, anterior temporal, and limbic regions with PFC regions and are expected to run or terminate in close proximity to UF and AmF. ICa was seeded in the ventral section of the anterior limb of the internal capsule passing between the caudate and pallidum, superior to the anterior commissure, and inferior and lateral to the caudate, in a position similar to that used by Jbabdi et al. (2013). The EmC seed mask was drawn in the sheet of white matter between the putamen and insula. The seed voxels were placed as laterally as possible within this sheet to favor fibers of the extreme over the external capsule, although due to the resolution of diffusion imaging it is not possible to fully ensure that no fibers of external capsule were included in the reconstructed EmC (Mars et al., 2016a). EmC seed voxels were placed just superior to the UF, in order to avoid including fibers belonging to this latter tract. The CB seed mask was drawn in a coronal plane capturing the WM dorsal and medial to the corpus callosum, within the cingulate gyrus.

To constrain the tractography algorithm, exclusion masks were drawn in both species to exclude false positive results in areas of high crossing fibers as follows: (1) within the basal ganglia to avoid picking spurious subcortical tracts (except for ICa tracking); (2) posteriorly to the seeds to prevent the projections from running backwards as the frontal cortical connections were the focus of this study; (3) an axial slice at the level of the superior temporal gyrus to prevent tracts from running in a ventral direction in an unconstrained manner (except for UF, CB, and AmF); (4) on the axial-coronal slices cutting across the thalamus, basal ganglia, and corpus callosum to exclude these subcortical and callosal projections; (5) in the dorsal cingulate cortex to avoid leakage of tracts to nearby bundles as a result of high curvature of the tracts (except for CB tracking); (6) an L-shaped coronal mask from the paracingulate cortex and the inferior frontal sulcus to the vertex to exclude tracking in the superior longitudinal fascicle; and (7) the opposite hemisphere to only track ipsilateral tracts. An exclusion mask of the CSF was used in each subject to prevent fibers from jumping across sulci during tracking. Streamlines encountering any of the exclusion masks were excluded from the tractography results. A coronal waypoint section was drawn in the frontal lobe at the level of the caudal genu of the corpus callosum to ensure that the fibers emanating from the seeds were projecting to the prefrontal cortex; this prevented leakage of the EmC into the corticospinal tract.

Probabilistic tractography was performed in each individual’s diffusion space. All seeding and tracking parameters were kept constant across tracts and only adjusted between species to deal with differences in brain size and data resolution. Human tractography parameters were: maximum of 3,200 steps per sample; 10,000 samples; step size of 0.25 mm; curvature threshold of 0.2. Macaque tractography parameters were: maximum 3,200 steps per sample; 10,000 samples; step size of 0.1 mm; curvature threshold of 0.2.

A visitation map or ‘tractogram’ was constructed for each individual in order to allow comparison of these maps between tracts, subjects, and species. Each tractogram was log-transformed to account for the exponential decrease of visitation probability with distance and normalised by dividing each voxel’s value by the 75th percentile value across the tractogram, thereby removing potential bias of differences in numbers of streamlines between tracts and across species. In both human and macaques, the focus of the investigation was on the tracts’ connections with prefrontal cortex. Prefrontal projections of each tract were averaged in a species-specific group-template tractogram. For the purposes of visualization only, the normalized tractograms were subsequently thresholded with minimum and maximum values equal to 0.5 and 2, respectively.

### Connectivity fingerprints

The differential connections of the reconstructed tracts with areas in prefrontal cortex were quantified by creating so-called connectivity fingerprints (Mars et al., 2018; Passingham et al., 2002) of each tract with target regions in frontal cortex. Such fingerprints can be used to compare tracts’ frontal projections with one another, but also to compare the tracts across species. Human and macaque frontal regions of interest (ROIs) were created based on previously published coordinates (Neubert et al., 2015, 2014; Sallet et al., 2013) or drawn by hand on the basis of anatomical atlases (Mai et al., 2015; Paxinos et al., 2000; Petrides et al., 2012; Yeterian et al., 2012) (Figure S2). Each coordinate was projected onto a subject-specific cortical surface mesh with 10,000 vertices (10k) in the human and to a template 10k surface in the macaque. Here, on the surface representation, these coordinates were expanded to circular ROIs following the cortical contours, with a 10 mm radius in humans and a 3 mm radius in macaques. The resulting ROIs were then projected back to each subject’s native brain image in volumetric space. The creation of ROIs on the surface rather than in volumetric space allows us to account for individual idiosyncrasies in sulci and gyri anatomy, ensuring that we have truly cortical rather than spherical ROIs. The white matter/gray matter border of each ROI was then extracted and used as target region in the connectivity fingerprint. This approach allows to maximize the estimation of streamlines projecting to each region while minimizing the reduction on signal consequent to tissue-related poor anisotropic signal characteristic of ROIs created in gray matter only. The fingerprints are then constructed by counting for each tract the average number of streamlines hitting the voxels of each ROI. To allow comparison across tracts and species, these values were normalized by dividing each tract-fingerprint by its maximum values across the selected ROIs. Connectivity fingerprints can then be compared by calculating the Manhattan distance between them (Mars et al., 2016b).

## Data availability

Results and recipes for tracking the tracts described here will be made available from the lab’s website (www.neuroecologylab.org). Raw and preprocessed human data are available from the Human Connectome Project (www.humanconnectome.org).

## RESULTS

Our goal was to reconstruct the amgydalofugal and uncinate pathways connecting the amygdala and anterior temporal cortex with the frontal cortex. In particular, we were interested in assessing the course of these pathways towards the frontal cortex and their frontal termination points. Given the difficulty of reconstructing these pathways, we employed the following strategy. First, we reconstructed the pathways in the macaque monkey brain, so that they can be compared to known tracer results (Figure 1A-C). This demonstrated whether our tractography techniques are valid for these pathways. Then, we applied the same tracking strategy to the human, using the seeding and masking procedures applied successfully in the macaque (Figure 1D-F). Finally, we compared the tracts’ projections directly and across species and with respect to a number of control tracts (Figure 3).

**Figure 1.**
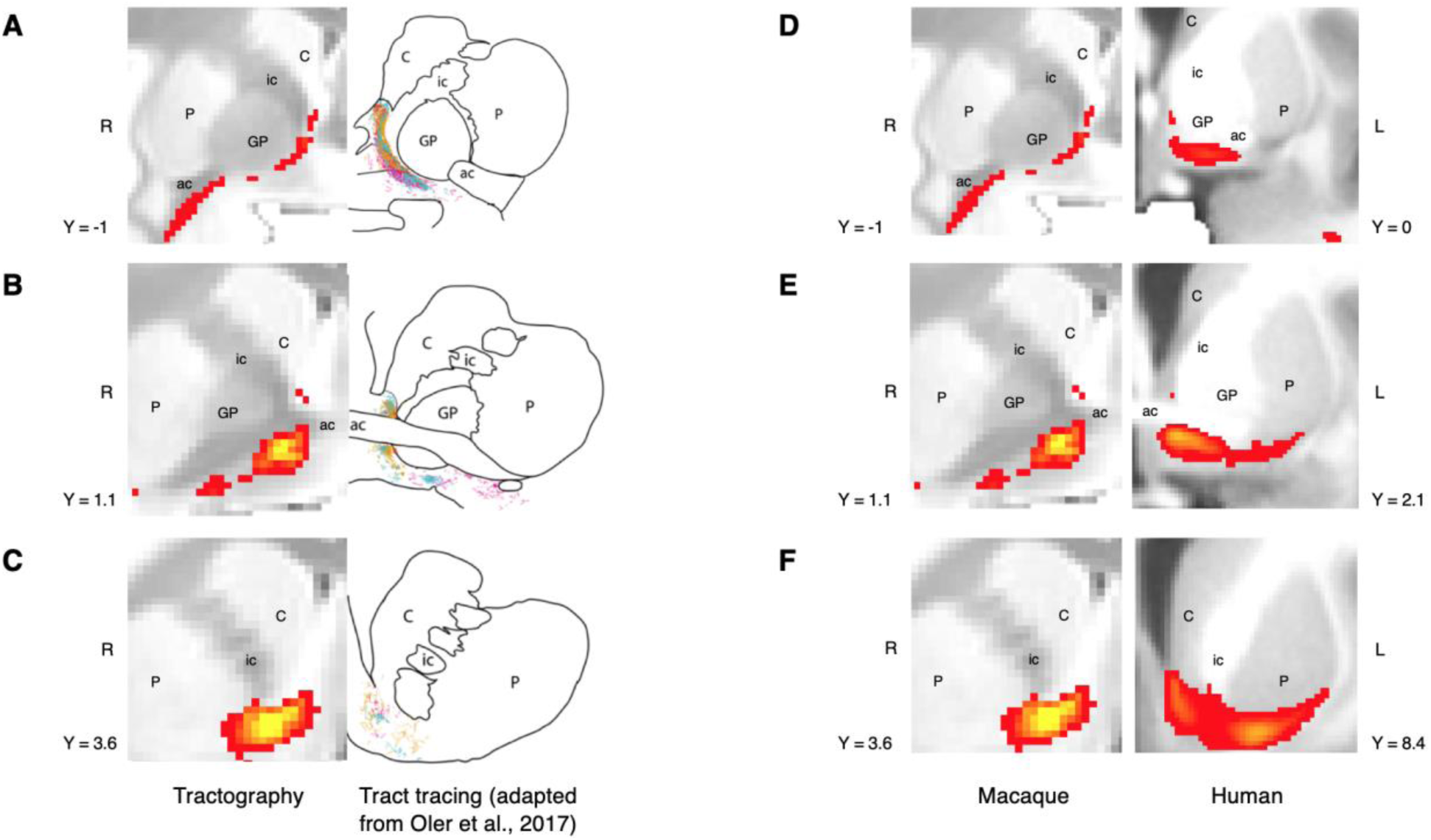
Histology-informed approach for the reconstruction of the frontal limb of the ventral amygdalofugal pathway. **(A-C)** Coronal sections depicting side-by-side the reconstruction of the frontal limb of the amygdalofugal pathway (AmF) in the macaque brain using probabilistic tractography on diffusion MRI data (left panels) and published reconstructions of the tract using radio-labeled fibers identified using tract tracing injections in macaque (right panels; adapted from (Oler et al., 2017). The AmF tractography reconstruction showed marked correspondence in anatomy and course of the fibers with the histological tracing data. AmF fibers leave the amygdala caudal to the anterior commissure (ac) and turn medially towards the medial wall of the hemisphere (A) and rostrally towards the frontal lobe by coursing ventrally to the ventromedial striatum (B-C). **(D-F)** The histology-informed reconstruction of the AmF was then used to guide the tractographic algorithm in the human brain, including the seed, exclusion and waypoint masks were used in both species (see Methods for details on the masks). The human AmF (D-F, right panels) follows a course matching the results observed in the macaque brain (A-C) with fibers projecting both medially (D) and rostrally (E-F). Note the macaque T1-weighted image (D-E, left panels) contrast-to-noise ratio is inverted because the brain sample was scanned ex-vivo, conversely to the human in-vivo samples (D-E, right panels). Ex-vivo acquisition of T1-weighted images are characterized by longer relaxation times of brain tissue which causes the white/gray matter contrast to be inverted compared to standard in-vivo acquisitions. T1-weighted image are shown in radiological convention. ac, anterior commissure; C, caudate nucleus; GP, globus pallidus; ic, internal capsule; P, putamen; L, left hemisphere; R, right hemisphere. *See also Supplementary Figures S1 and S3*.

### Amygdalofugal pathway in the macaque monkey brain

The frontal projections of the AmF are represented by a thin bundle of fibers running in the WM adjacent to medial subcortical and cortical brain structures (Figure 2A-G, S3A-C). In a posterior section of the macaque brain, fibers running in the medially projecting limb of the AmF pathway initially left the amygdala coursing dorsally to then turn in the direction of the medial longitudinal fissure at the level of the posterior anterior commissure (AC) and sublenticular extended amygdala area (SLEA) (Figure 1A-B, 2A, S3A-B). Here, a first bundle of AmF fibers ran in the white matter territory in between the nucleus basalis of Meynert, lentiform nucleus, ventral pallidum, and the piriform cortex (Figure S3A-B). The medial AmF fibers are known to separate into different sets of projections (Lehman et al., 2011; Nauta, 1961; Novotny, 1977; Oler et al., 2017) that extend ventrally to innervate the anterior hypothalamic areas at the level of the paraventricular-supraoptic nuclei and dorsally to connect the amygdala with the thalamus, the BNST, and septum. All the fibers were visible in our data to leave the seed point, but the frontal waypoint mask means the pathway to the frontal cortex, which is the focus of the current investigation, is most prominent (Figure 2A,B, S3A,B). This second, rostrally projecting bundle of the AmF left the substantia innominata by turning rostrally throughout the ventral pallidum and sub-commisural white matter to continue in a rostral direction in the region ventral to the ventral striatum/nucleus accumbens (VS/NAc; Figure 1C, F, 2C-G). From tract tracing, we know that some fibers connecting ventral PFC and amygdala are intermingled with the much more prominent anterior commissure (Lehman, et al. 2011). The AC is very prominent in the principal fiber direction of diffusion MRI data, but nevertheless some weak AmF projections overlapping with AC were evident in our tractography data (Figure 1B,1E, 2A, S3B).

**Figure 2.**
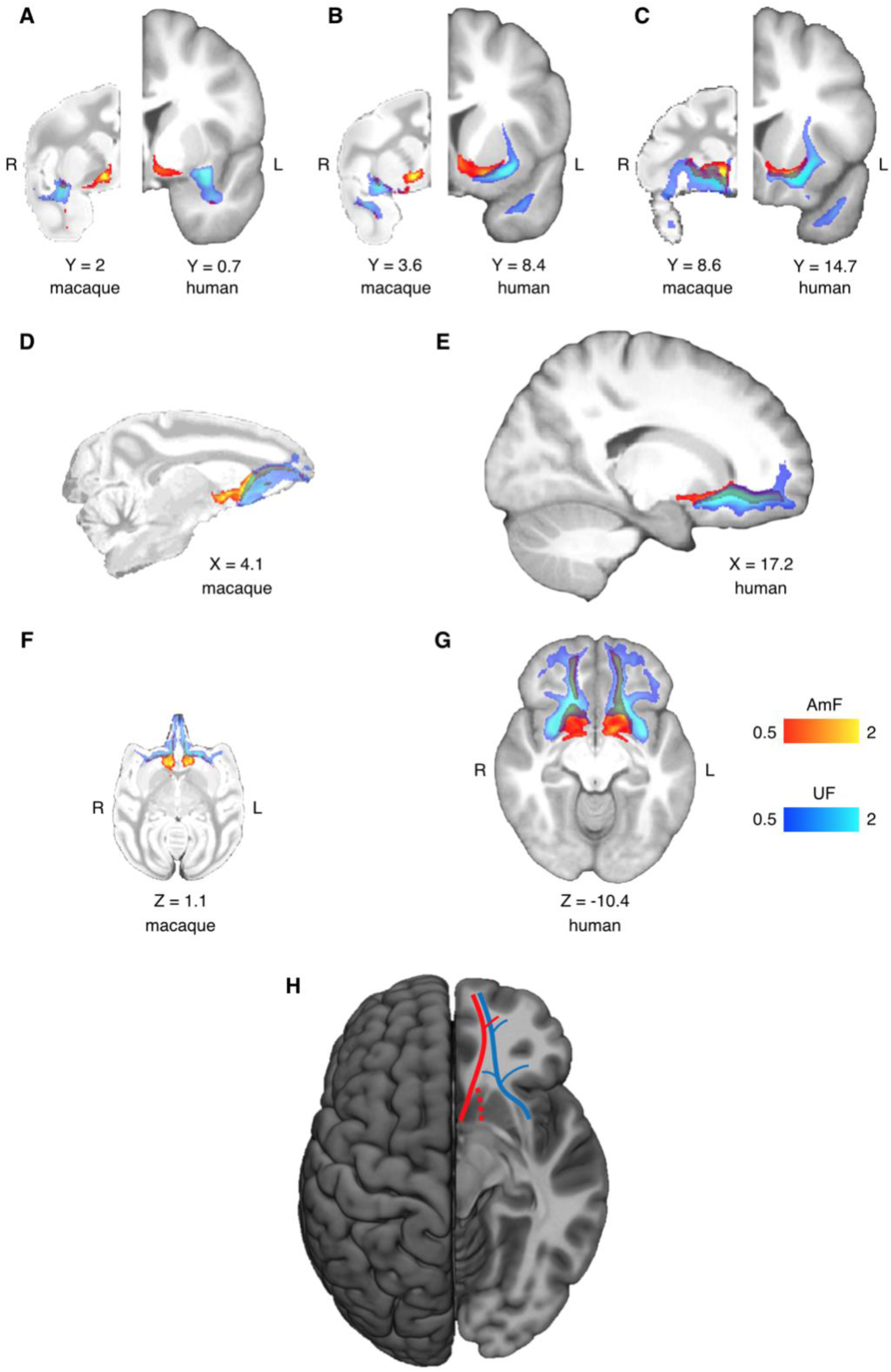
Anatomy and organization of the Amygdalofugal Pathway and Uncinate Fascicle in the basal ganglia and frontal lobe of the macaque monkey and human brain. **(A-C)** amygdalofugal pathway (AmF, red) and uncinate fascicle (UF, blue) are displayed on three coronal brain slices. AmF and UF show the same medial-lateral organization in both macaque (left) and human (right) brains. **(D-E)** Sagittal view of the macaque (D) and human (E) brain displaying the preservation of the dorsal-ventral and medial-lateral organization of the two tracts across both species in caudal subcortical regions of the brain but their merge in caudal orbitofrontal cortex. **(F-G**) The medial-lateral organization of the two tracts is shown on axial sections of macaque (F) and human (G) brains. The AmF follows a course medial to the UF with its fibers wrapping the full extent of the ventral striatum and nucleus accumbens in both macaques (F) and humans (G). Here, the main body of the UF runs instead more laterally, primarily in the WM adjacent to the ventral-lateral striatum and the insular cortex. The tracts merge into an intermingled connectional system in the posterior OFC (area 13) and for the full extent of the ventromedial frontal cortex. Macaque coordinates are in F99 space and human coordinates are in MNI space. **(H)** Schematic template representing the organization of AmF (red) and UF (blue) in the ventral frontal in the human brain. Dashed line (red) shows a bundle subcortical AmF fibers running underneath and through the ventral striatum. *See also Supplementary Figures S1 and S3*.

It must be noted that over its course in the basal forebrain and ventral to the striatum, the AmF is likely to interdigitate with other connectional systems including the rostral fibers of the thalamic projections, the medial forebrain bundle, the diagonal band of Broca and the dopaminergic fibers connecting the ventral tegmental area with prefrontal cortex. Eventually, the anterior projections of the AmF reached the orbital periallocortex on the medial wall of the hemisphere and the PFC by entering the subgenual anterior cingulate cortex (sACC or Brodmann area 25; Figure 2C-D,F, S3C).

Near the sACC, the AmF split again into two sub-sections. The first sub-section runs alongside the cingulum bundle (CB) travelling dorsally towards the anterior cingulate sulcus and perigenual anterior cingulate gyrus. The second branch traveled from the sACC towards the frontopolar cortex passing subjacent to medial orbitofrontal cortex (OFC) between the gyrus rectus and the medial orbital gyrus (Figure 2D and 2F). Because diffusion MRI cannot discern between monosynaptic and polysynaptic connections, the identified fibers could represent either type of projection.

### Amygdalofugal pathway in the human brain

As in the macaque, in the human brain the AmF pathway ran adjacent to the lateral and basal nuclei of the amygdala and curves in the more dorsal WM within the substantia innominate and SLEA bordering the central and medial nuclei where additional fibers entered the main body of the tract (Figure 1D-E, 2A, S3A-B). Here, the AmF extends subcortically along the medial wall and gives rise to two limbic bundles. Consistent with the macaque, the first set of AmF fibers takes a medially projecting path that follows a trajectory resembling a reversed S-shaped coursing from the caudal amygdala towards peri-ventricular regions located both superiorly and inferiorly to the AC and from here it ran towards the bed nuclei of stria terminalis, hypothalamus and thalamus. A second portion of the ventral AmF pathway left the substantia innominata heading rostrally close to the SLEA by interdigitating with AC fibers or running through the sub-striatal WM surrounding the VS/NAc (Figure 2A-B, S3A-B).

The human AmF entered the PFC predominantly through strong innervation of the sACC although a smaller contingent of fibers connect with caudal OFC. In ventral PFC, the main body of the AmF kept running along the medial bank of the cortex in a position adjacent to ventromedial PFC (vmPFC) in the gyrus rectus and medial OFC (Figure 2C; Figure 2E; Figure 2G and Figure S3C). Recently, Neubert et al. (2014) suggested that the frontopolar cortex (FPC) in the human brain is composed of functionally specialized medial and lateral sub-regions to a greater extent than in the macaque. The AmF projections kept running close to the medial wall in the FPC but fewer projections also extend laterally to reach the lateral FPC (Figure 2E; Figure 2G).

The architecture of the human AmF closely resembled that of the macaque brain, suggesting that this tract is evolutionarily preserved; both its medial and anterior branches are very similar in the human and macaque monkey brain (Figure 2A-G) despite the limited white matter in the latter species (Schoenemann et al., 2005). Minor differences arise when we look at FPC; in the human few AmF fibers headed towards the lateral FPC and the majority of the tract innervates medial FPC (Figure 2E,G).

### Uncinate fascicle in the macaque monkey brain

The UF is a hook-shaped tract containing fibers interconnecting the temporopolar cortex and amygdala with orbital gyri and ventrolateral PFC in a bidirectional manner. On a ventral section, the macaque UF extended along the lateral surface of the temporal lobe where it made a dorsal C-shaped curve through the superior temporal gyrus of the temporal lobe, coursing medially to the Sylvian fissure (Figure 2F and Figure S3D). Although in this region the principal fiber direction runs in a superior-inferior plane, some fibers branch along a medial-lateral axis to connect with the amygdala and carry interconnections between amygdala, the temporal lobe, and lateral frontal cortex (Figure 2A,F, S3E). Other fibers also leave the main body of the tract and run caudally to innervate the parahippocampal gyrus and rhinal cortex (Figure 2F, S3D-E). At the level of the limen insulae, UF fibers run in a white matter region ventrally to the claustrum, ventrolaterally to the putamen and globus pallidus, and medially to the insular cortex. Here, the most dorsal fibers of the UF intermingled with the ventral axons of the EmC as well as connecting with ventrolateral PFC (Figure 2A-C, S3D-F).

In monkey PFC, the UF strongly innervated the caudolateral OFC, the opercular and inferior frontal areas although a sub-section of fibers reached the subgenual cingulate area and the medial bank of posterior OFC (Figure 2C-D and Figure S3F). Here, the more medial UF fibers joined with the AmF and CB fibers to innervate ACC (Figure 2D). Overall, the main body of the UF in ventral PFC stretches throughout lateral, central, and medial orbital gyri giving rise to a complex network in OFC which extends into the frontopolar cortex (Figure 2F).

### Uncinate fascicle in the human brain

The human UF anatomy has a preserved C-shaped structure connecting rostral and medial-temporal regions with the ventrolateral PFC (Figure 2E, S3D). Similarly to macaque temporal UF, it is anatomically organized predominantly along a dorsal-ventral gradient but with lateral fibers connecting it with the amygdalar nuclei and parahippocampal and rhinal areas (Figure 2G, S3D-E).

In the intersection between prefrontal and temporal cortices, it innervates the insular cortex and regions in the caudal territory of the inferior frontal gyrus (Figure 2B-C,G, S3D-F). More medially, human UF encompassed the lateral ventral striatum and kept a predominant lateral course as it enters the ventral PFC (Figure 2A-C, S3D-F), whereas a bundle of UF fibers join the cingulate AmF and CB (Figure 2E). Here, the main body of the UF kept running in a lateral position sending widespread innervation to all the ventrolateral PFC. Furthermore, a sub-section of fibers project medially, partly to sACC and more prominently to posterior OFC (Figure 2C,G, S3F).

In the anterior territory of ventral PFC the human UF innervated both medial and lateral orbital areas in a widespread fashion. Strong projections reach the lateral OFC and adjacent cortical areas located on the ventral surface of the brain. Another laterally projecting bundle leaves the main body of the UF more rostrally and innervates the lateral FPC, whereas medial FPC is innervated by the more medially coursing UF fibers (Figure 2E,G, S3F).

Again, as for the AmF pathway, the anatomy of the UF as well as its interdigitation with AmF in the human brain is highly conserved in monkey and human brains (Figure 2A-G and Figure S3D-F). There may be differences in FPC organization between humans and macaques (Neubert et al., 2015) and it was notable that the UF innervated medial and lateral FPC in the human brain in equal proportion (Figure 2G).

### Comparison of the two fiber bundles

The main bodies of the human AmF and UF are anatomically segregated and constitute two anatomically distinct connectional systems in temporal, limbic, and striatal WM. Alongside the dorsolateral amygdala, AmF interdigitates with more lateral temporo-frontal UF axons. At this level, it is possible to identify in both species a medial-lateral organization of the two tracts with the former leaving the amygdala more caudally and the latter’s fibers projecting to and from this subcortical structure in a more anterior position (Figure 2).

In the basal ganglia (BG), the two tracts maintain a clear and evolutionary preserved organization along a medial-lateral gradient, but also start to develop a dorsal-ventral relationship (Figure 2A-G). As previously described, in this region the AmF courses medial to the ventral striatum, just underneath the anterior limb of the ICa and medial to the ventromedial striatum, specifically inferiorly to the internal and external globus pallidus, the ventral head of the caudate and the medial bank of the putamen (Figure 2A-G and Figure S3A-B). Conversely, the UF approaches the BG more laterally. A medial portion of UF fibers interdigitates with lateral AmF axons and courses ventral to the AmF in the white matter surrounding the putamen. In contrast to the more ventromedially positioned course, UF fibers run alongside the full ventrolateral extent of the BG. In the WM lateral to the putamen and ventral to the claustrum, its fibers interlace with the ventral axons of the EmC (Figure 2A-G and Figure S3E-F).

The relative mediolateral positioning of the AmF and UF extends more rostrally into the WM adjacent to the sACC (Figure 2C and Figure 2F-G); a major component of the AmF lies adjacent to sACC (Figure S3C) while only a medial bundle from the UF lies adjacent to sACC (Figure 2C and Figure S3F). The relative positioning of the two pathways then begins to partially rotate with the UF moving to a location ventral to the AmF as the anatomical distance between the two pathways reduces in the subgenual WM. In the white matter adjacent to infralimbic cortex the segregation of the two tracts begins to fade as the dorsomedial UF fibers approach the ventrolateral AmF axons (Figure 2C and 2F-G).

Although the AmF and UF are clearly distinct entities in more posterior parts of the brain their clear separation disappears in the posterior OFC. Here, the UF and AmF merge into a single connectional system. This anatomical organization is present and conserved across both human and macaques in relation to several homologous landmarks in both species (Figure D-G; Figure S3C and Figure S3F).

### Prefrontal connectivity fingerprints

We have described the distinct courses of the AmF and UF from the vicinity of the temporal cortex and amygdala towards the frontal cortex. To quantify the projections of the tracts to frontal areas, we can describe their connectivity fingerprints with frontal regions of interest (Mars et al., 2018). This allows a qualitative and quantitative comparison of the areal connectivity of AmF and UF. To examine the specificity of the results, we also compared their connectivity fingerprints to those of three control tracts connecting nearby systems: the anterior limb of the internal capsule (ICa), the extreme capsule (EmC), and the cingulum bundle (CB) (Figure 3).

**Figure 3.**
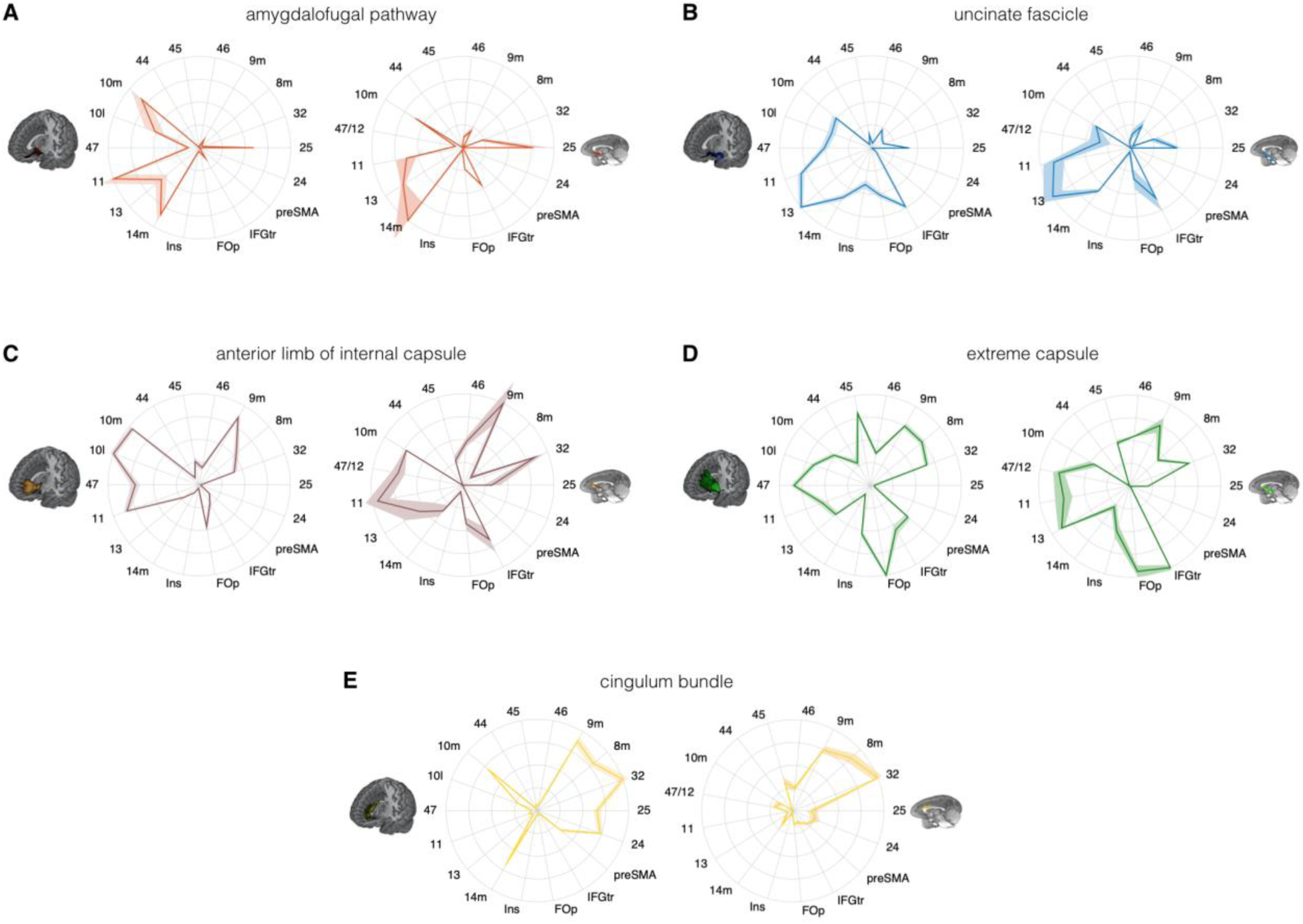
Comparison of connectivity fingerprints representing the projections of the amygdalofugal pathway, uncinate fascicle and adjacent white matter tracts in the human (left) and macaque (right) frontal lobe. **(A)** The pattern of AmF projections is preserved across the two species and predominantly targets medial areas in subgenual, orbital, ventral, and polar PFC. (**B**) Overall UF connections show high similarity between the two species but stronger projections to the opercular and insular cortex is observable in the human UF. (**C-E)** Macaque - human comparison of the frontal projections of three connectional systems located in the white matter dorsal to the AmF (internal capsule; C), dorsal to the UF (extreme capsule; D) or interacting with both tracts in the infralimbic and prelimbic cortex (cingulum bundle; E). 9m, dorsomedial PFC; 8m, caudal bank of the dorsomedial PFC; 32, anterior cingulate cortex; 25, subgenual cingulate cortex; 24, dorsal anterior cingulate cortex; preSMA, pre-supplementary motor area; IFGtr, inferior frontal gyrus, pars triangularis; FOp, frontal operculum; Ins, insular cortex; 14m, ventromedial PFC; 13, caudal orbitofrontal cortex; 11, rostral orbitofrontal cortex; 47 (human) or 47/12 (macaque), lateral orbitofrontal cortex (also named lateral convexity of PFC, Murray and Rudebeck, 2018; Rudebeck and Murray, 2014); 10l, lateral frontopolar cortex; 10m, medial frontopolar cortex; 44, caudal bank of the ventrolateral PFC; 45, rostral bank of the ventrolateral PFC. Shades represent standard error of the mean (SEM).

Comparing the connectivity fingerprints of AmF and UF show that in both species, a medial-lateral organization in projection strength between the two tracts is evident (Figure 3A-B). While both pathways were primarily interconnected with ventral frontal cortex in both species, AmF was more likely to reach medial areas such as 25, 14m, 11 and 10m (Figure 3A), whereas UF primarily reached more lateral areas such as the inferior frontal gyrus, insular cortex, frontal operculum, area 47 (or the macaque homolog 47/12), and area 45 (Figure 3B). However, UF fibers in the rostral ventral PFC were also connected with more medial OFC (area 11) and FPC (10m). Some species-specific differences were also evident. Human AmF projects more prominently to area 11 and 10m, whereas macaque AmF preferentially innervates area 32 and the caudal vmPFC area 14m (Figure 3A). Inter-species differences in cingulate projections may partly be the results of species-specific idiosyncrasies in neuroanatomical geometries, especially when diffusion MRI is used to reconstruct tracts running in the WM adjacent to brain structure presenting different degrees of curvatures in different species. UF also exhibited some contrasting inter-species differences. Cortical projections to lateral orbital, ventrolateral, insular, and opercular areas were stronger in the human compared to macaque brain, suggesting a more widespread pattern of connectivity in human frontal cortex (Figure 3B). This may reflect prefrontal cortical expansion in humans compared with other old-world primates (Smaers et al., 2017). As discussed above, in the most anterior part of prefrontal cortex, AmF and UF tend to intermingle and it is therefore not surprising to see that both reach the frontal pole (area 10m), including the human lateral frontal pole (area 10l) for which the homology in the macaque monkey brain is debated (Neubert et al., 2015).

The connectivity fingerprints of the control tracts were markedly distinct from those of both AmF and UF (Figure 3C-E). The anterior limb of the internal capsule carries fibers originating and terminating in the thalamus and brainstem (Lehman et al., 2011). Its frontal projections reached more dorsomedial regions than AmF and UF, including areas 32, 8m, and 9m (Figure 3C). EmC reached even more widespread areas of lateral and dorsal prefrontal cortex, including areas 45, 8m, 9m, and 32 (Figure 3D). The cingulum bundle revealed a projection pattern primarily reaching medial areas (Figure 3E). Interestingly, similarly to the UF, some branches of these control tracts exhibited a more widespread pattern of connectivity in the human compared to macaque brain. Indeed, human EmC projected more caudally in dorsal PFC, reaching the preSMA (Figure 3D). CB fibers, instead, showed stronger projections in frontopolar, orbital, subgenual and midcingulate cortex compared to the macaque (Figure 3E).

The connectivity fingerprints can be used to quantitatively compare the connectivity profiles across species. We can create a dissimilarity matrix between species by calculating the Manhattan distance for each possible pair of tracts. This measure indicates how different two connectivity fingerprints are, whereby the best matching pairs are defined as the set of fingerprints that have the smallest distance values (Mars et al., 2016b). When comparing the fingerprints (averaged across the two hemispheres) between species, we see that the smallest differences for each tract are on the diagonal of the distance matrix (Figure 4), indicating that each tract has a connectivity fingerprint most similar to its counterpart in the other species. AmF and UF are each other’s second best choice, but match best with their counterpart. This similarity is not unexpected given close proximity in ventral PFC and considering the strong interconnection and reciprocal horizontal projections of OFC regions, yet despite these correspondences the dissimilarity matrix confirms the two pathways are distinguishable.

**Figure 4.**
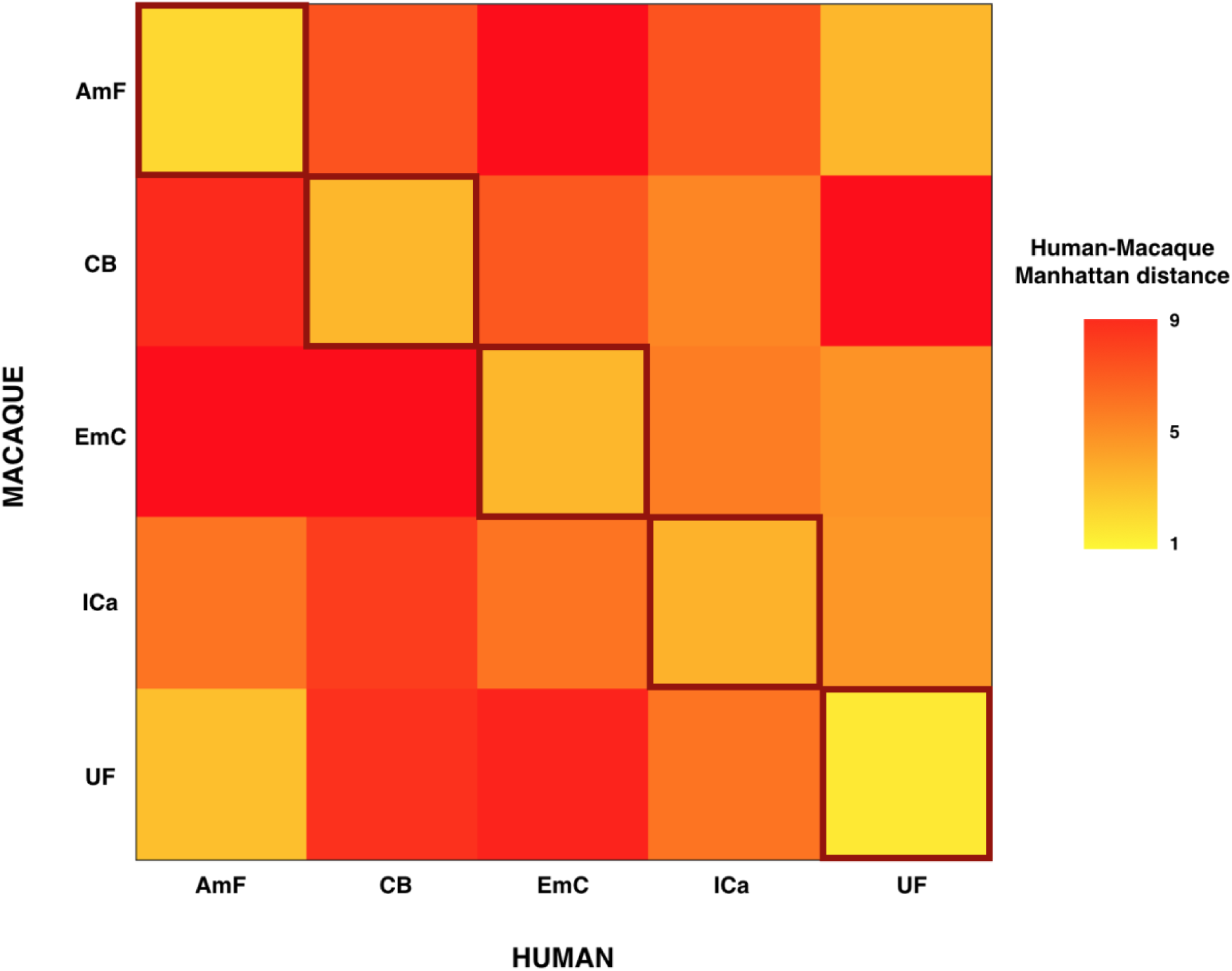
Quantitative comparison of the connectivity fingerprints of the five reconstructed frontal tracts in the human and monkey brains. Dissimilarity matrix of the frontal projections of the tracts of interest across species. Human tracts are listed on the horizontal axis; macaque tracts are listed on the vertical axis. Pair-wise dissimilarity is defined as the “Manhattan distance” between each pair of pathways across the two species. The greatest similarity was found when comparing each tract with its homologue in the other species (bright diagonal in the dissimilarity matrix). AmF, ventral amygdalofugal pathway; CB, cingulum bundle; ICa, anterior limb of the internal capsule; EmC, extreme capsule; UF, uncinate fascicle.

## DISCUSSION

In this study we identify and define two closely related but distinct connectional systems linking amygdala and PFC using diffusion MRI tractography in primates. The structure, course, and projections of these tracts in the macaque monkey closely match previous tracer injection studies (Fudge et al., 2002; Fudge and Haber, 2001; Kunishio and Haber, 1994; Lehman et al., 2011). Moreover, we showed that both anatomy and connectional profiles of these two tracts are preserved across humans and monkeys. We demonstrated that both the AmF and UF run separately and in parallel when exiting the medial temporal lobe but then merge to form a single fiber complex in the ventral PFC.

Tract-tracing studies provide important information about where specific axons in fiber bundles terminate or originate in the brain (Petrides and Pandya, 2007). However, the technique is invasive and requires the sacrifice of the subject, limiting its use in human anatomy. Diffusion MRI tractography excels when tracing in homogenous white matter and can reconstruct many of the major white matter tracts. However, it can fail in areas in which fiber density is low or different fibers cross. These properties are exhibited by the porous heterogenous tissue in the basal forebrain that the amygdalofugal pathway traces through, as evidenced by invasive tracer injection and histology studies. It would be especially difficult to track in these circumstances using algorithms that do not utilize the full probability distribution of the fiber directions. Perhaps because of these difficulties, previous studies using tractography to investigate the limbic system might have overlooked or undervalued the frontal projections of the AmF (Catani et al., 2013; Croxson et al., 2005; Kamali et al., 2016). In contrast, to overcome these challenges, we employed a strategy of first performing probabilistic tractography based on crossing-fiber resolved diffusion models in the macaque monkey to develop a protocol that was able to reconstruct the AmF fibers as known from previous invasive tracer studies in that species (Oler et al., 2017). This strategy was successful. We were able to reconstruct the course of the tract and confirmed the AmF as primarily a subcortical limbic tract, but with frontal connection mostly extending to medial areas such as 25 and 14m. This pattern was similar in the human.

The division between AmF and UF connections maps onto the tracer-based division of the ventral PFC into a medial region and a more lateral/orbital region as originally suggested by Price and colleagues (Carmichael and Price, 1995; Ongür et al., 2003). Following the logic that the connections of an area are a crucial determinant of its function, this suggested a different functional role of these subdivisions with the medial network strongly connected with visceromotor areas and the lateral orbital network interconnected with multi-modal sensory inputs (Ongür and Price, 2000). This was confirmed by dissociations apparent from lesion studies with medial frontal lesions showing an impairment in the capacity to maintain sustained arousal when anticipating rewards (Rudebeck et al., 2014), but temporal and ventral lateral frontal lesions leading to an impairment in learning arbitrary associations between stimuli and outcomes or stimuli and actions (Browning and Gaffan, 2008; Eacott and Gaffan, 1992; Gutnikov et al., 1997). Our results show divergences in inputs to ventral PFC that supports this dissociation and demonstrate that it generalizes to the human brain. Fibers in AmF and UF may constitute the primary white matter scaffolding from which these two networks may arise.

It is important to note, however, that the dissociation between medial and lateral circuits is not complete, with the circuits joining up in anterior OFC and rostral prefrontal cortex. An integration between the two processing streams has been suggested both on anatomical and functional grounds. Anatomically, more rostral parts of prefrontal cortex are suggested to be more interconnected (Carmichael and Price, 1995). Functionally, such an integration would subserve learning and decision-making computations by allowing the integration of learned stimulus-outcome contingencies with the internal state of the organism. This is evident in the proposal of a “common currency” representation comprising the value of different objects irrespective of the physical properties (Montague and Berns, 2002; Padoa-Schioppa and Assad, 2008). The rostral interactions of AmF and UF in areas 11, 13 and 14m may therefore represent the anatomical support for such computations.

The importance of considering neural circuitry, extending beyond individual brain regions, is increasingly evident when studying the contributions of ventral PFC in emotion, learning, and decision-making. Combining controlled focal lesion and transient perturbation studies with whole-brain neuroimaging highlights the critical role of remote connections to the impact of local perturbations (Folloni et al., 2019; Froudist-Walsh et al., 2018; Verhagen et al., 2019). Moreover, it has recently been demonstrated that some lesion effects might be better explained by damage to connections as opposed to damage to single regions (Rudebeck et al., 2013; Thiebaut de Schotten and Foulon, 2018) and differences in connectivity have already been argued to play a crucial role in regional specialization in other brain areas (Krubitzer, 2007). Given the inverse problem in which different changes in brain activity can have multiple causes (O’Reilly et al., 2013), an understanding of connectional anatomy is therefore essential to understand these network effects. As such, this work can help to move from a view of ventral PFC as a constellation of isolated, “functionally specialized” regions to a more network-based approach, thereby helping to more appropriately describe the underlying neurobiology of psychiatric disorders and promote effective treatment options.

Conventionally, surgical interventions, including deep brain stimulation for depression, addition, and obsessive-compulsive disorders were primarily targeted at grey matter tissue. For instance, aspects of area 25 anatomy are associated with the presence of psychiatric disorders (Drevets et al., 1997; Murray et al., 2011). Indeed, chronic deep brain stimulation in this area improves chronic dysphoric conditions (Mayberg et al., 2005). However, with evidence accumulating that the most effective electrodes in deep brain stimulation are situated in or nearby white matter fiber tracts, rather than in grey matter proper, recent deep brain stimulation approaches are explicitly targeting white matter fiber structures (Baldermann et al., 2019; Lozano et al., 2019). Considering the effective electrical stimulation of area 25 (Johansen-Berg et al., 2008), this region is innervated and visited by the nearby AmF, suggesting the hypothesis that the physiological effects may be primarily ascribed to the AmF pathway and the regions it interconnects.

For this study we used high quality diffusion-weighted MRI data, obtained by scanning ex-vivo macaque samples for multiple hours, and by using in-vivo human diffusion MRI data with an acquisition duration of one hour. Both these datasets relied on unconventionally strong diffusion gradients, high number of diffusion directions, and high spatial resolution. However, the tractography recipes developed as part of this study have already been shown to be applicable and successful in human diffusion MRI data of more conventional quality. In fact, the approach and results achieved here, including the division between these two amygdala-prefrontal tracts, are behaviorally relevant in the context of social-emotional actions, advancing our understanding of the neurobiology of emotion regulation (Bramson et al., 2019 biorXiv).

To conclude, two principles underlie the organization of amygdala-PFC connectivity in the human brain. They are both evolutionarily conserved and shared with the macaque monkey. First a “connectional segregation principle” means that AmF and UF occupy a relative mediolateral position with respect to one another. Second, a “connectional integration principle” means that the tracts merge together as they enter and progress through the WM in the PFC to create an intricate network of projections from both cortical and subcortical regions. The two connectional systems described here may support distinct but synergic brain functions in PFC and this hypothesis should be investigated in future to understand how regions in the temporal lobe and PFC interact during affective, social, and cognitive tasks, how they could be disturbed in psychiatric disorders, and how this can be alleviated using targeted brain stimulation approaches.

## ACKNOWLEDGEMENTS

We would like to thank Matthew F.S. Rushworth and Richard E. Passingham for valuable discussions on the results. The work of D.F. is supported by the Wellcome Trust [UK Grant 105238/Z/14/Z]. The work of J.S. is supported by a Wellcome Trust Henry Dale Fellowship (105651/Z/14/Z). The work of L.V. is supported by the Wellcome Trust [WT100973AIA] and a Marie Curie Intra-European Fellowship within the European Union’s 7th Framework Programme [MC-IEF-623513]. The work of R.B.M. is supported by the Biotechnology and Biological Sciences Research Council (BBSRC) UK [BB/N019814/1] and the Netherlands Organization for Scientific Research NWO [452-13-015]. The Wellcome Centre for Integrative Neuroimaging is supported by core funding from the Wellcome Trust [203139/Z/16/Z]. NRS and AAK were supported by Cancer Research UK [grant C5255/A15935].

## SUPPLEMENTARY MATERIALS

**Figure S1.**
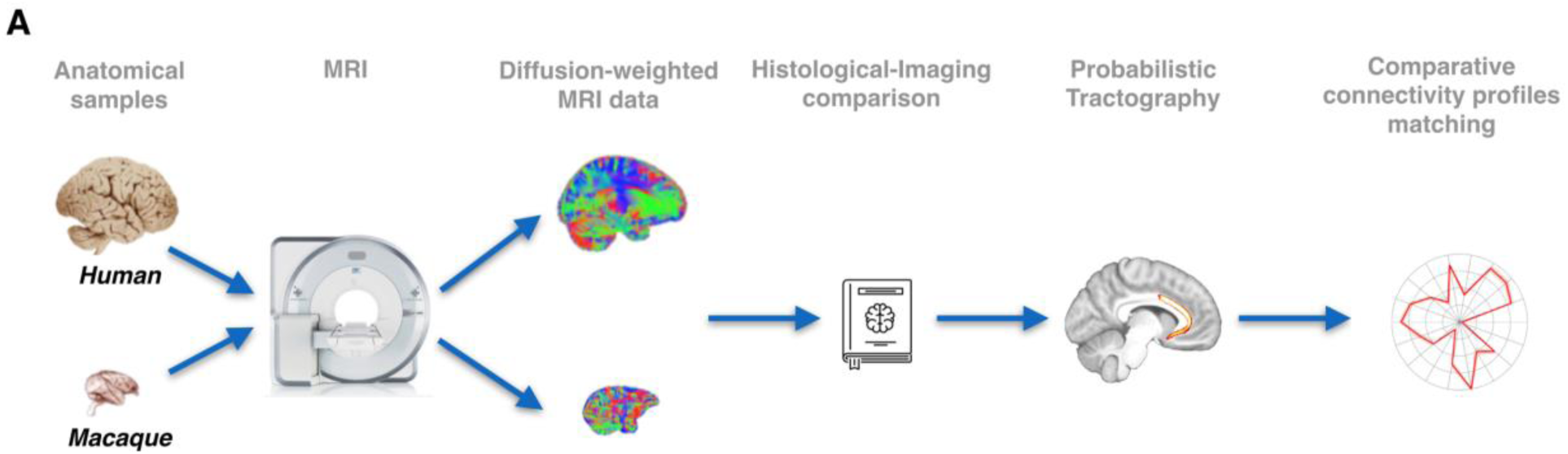
Methodological pipeline and white matter seed masks used to reconstruct each pathway. **(A)** We combined high-quality, high-resolution diffusion imaging data from in-vivo human (n=24) and post-mortem monkey (Macaca mulatta; n=3) with probabilistic tractography analyses to reconstruct the course and connectivity pattern of two tracts connecting amygdala and anterior temporal cortex with the frontal lobe: the ventral amygdalofugal pathway and uncinate fasciculus. For comparison, tractographic analyses were also run on three other limbic-PFC tracts: the anterior limb of the internal capsule, the extreme capsule and the cingulum bundle. Anatomical reconstruction of the 5 tracts was made by reference to previously published atlases (Nieuwenhuys et al., 2008; Paxinos et al., 2000), tract-tracing studies (Fudge et al., 2002; Oler et al., 2017; Schmahmann and Pandya, 2006) and diffusion tractography studies (Catani et al., 2013; Croxson et al., 2005). It was aimed to keep all seeding and tracking parameters constant across tracts and species. To constrain the tractography algorithm, *a priori* anatomical priors were used for the analyses and homologous tract-specific exclusion masks in both species were used in order to obtain high fidelity results.

**Figure S2.**
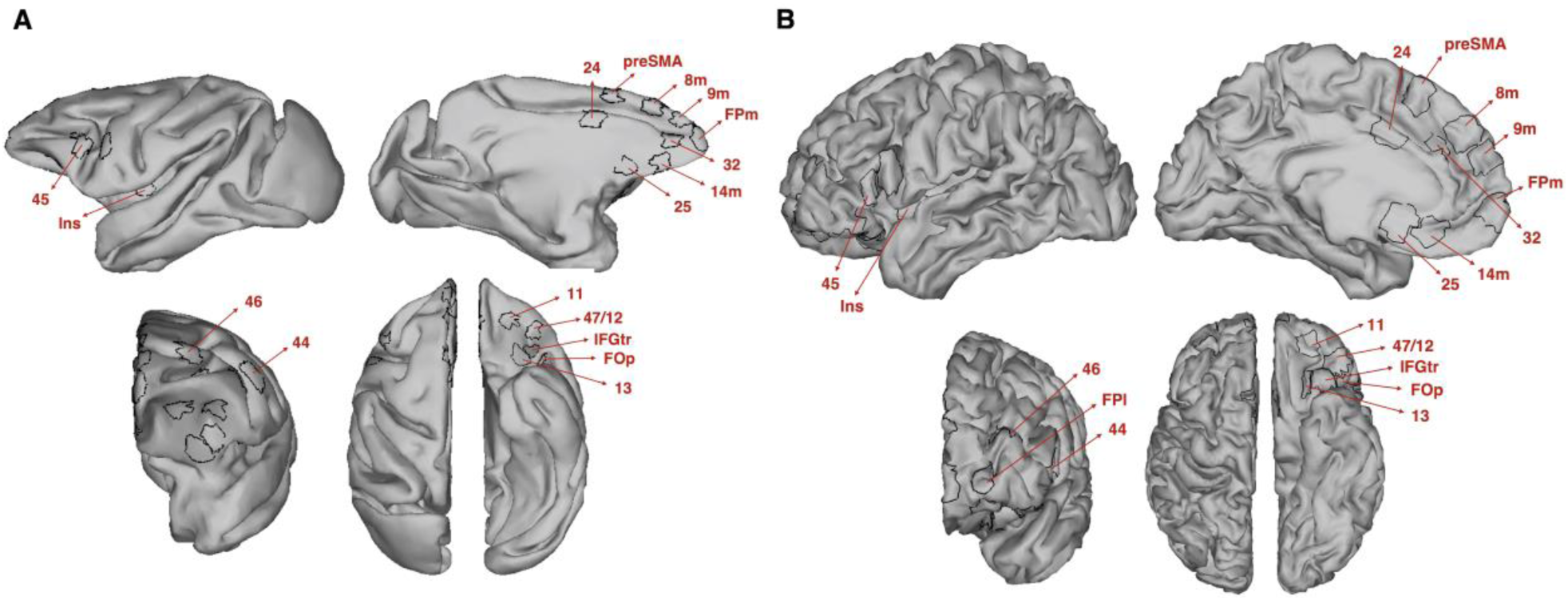
Target regions of interest used to estimate frontal connectivity fingerprint for each tract. **(A-B)** Regions of interest (ROI) were created on the cortical surface of the macaque (A; F99 space) and human (B; MNI space) brain. Each ROI was then projected to the volumetric gray matter / white matter border and used for the connectivity fingerprint analyses. All ROI labels as in figure 3.

**Figure S3.**
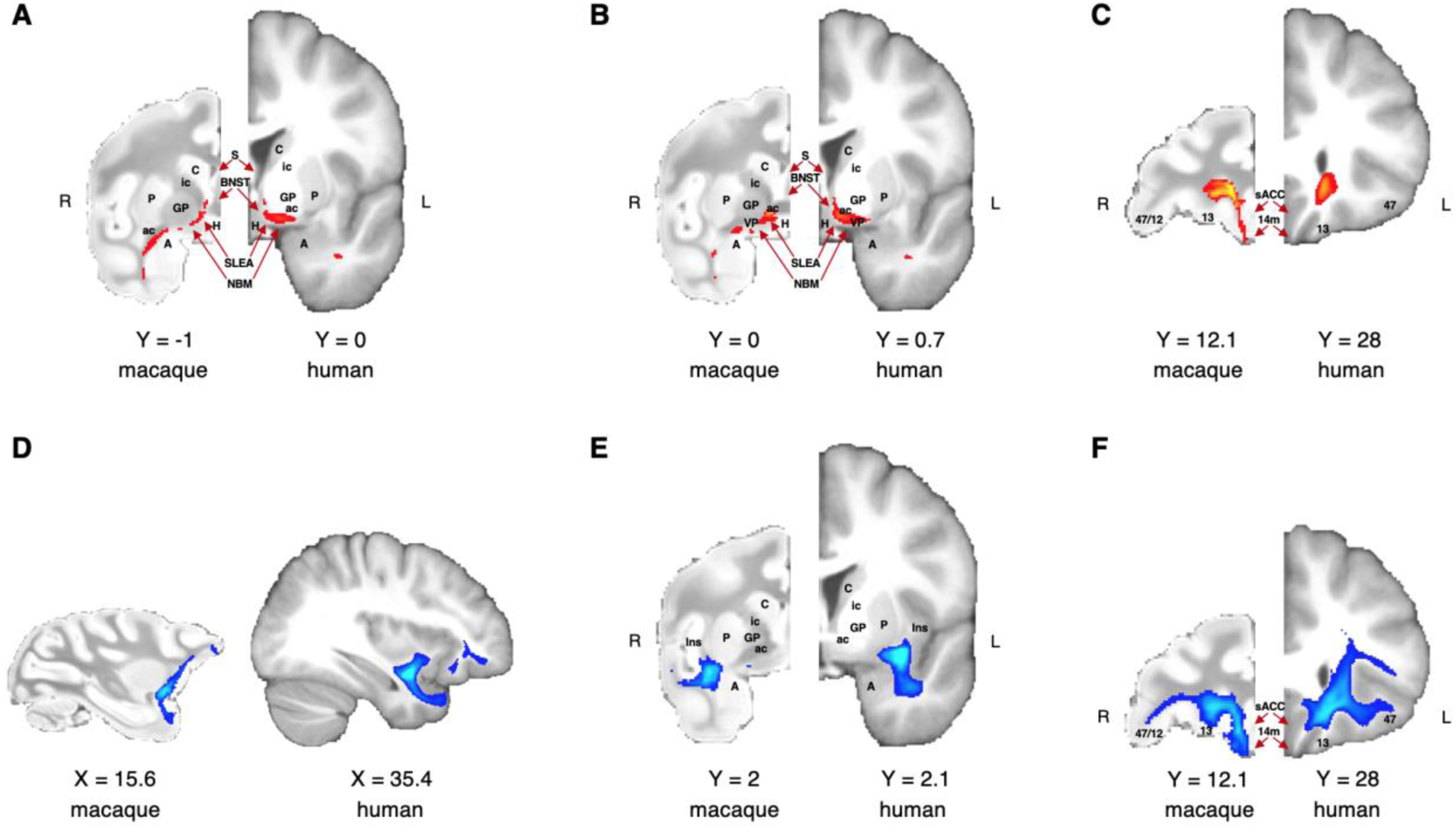
Comparison of the anatomical architecture of the ventral amygdalofugal pathway and uncinate fascicle in macaque monkey and human brains. **(A-C)** Anatomy of macaque and human amygdalofugal pathway on three coronal slices showed from caudal (A) to rostral (C). **(C-F)** Anatomy of macaque a human uncinate fascicle in a sagittal slice and in two coronal slices showed from caudal (E) to rostral (F). Macaque coordinates are in F99 space and human coordinates are in MNI space. A, amygdala; BNST, bed nucleus of stria terminalis; H, hypothalamus; Ins, insular cortex; NBM, nucleus basalis of Meynert; sACC, subegnual anterior cingulate cortex (Brodmann area 25); S, septum; SLEA, sublenticular extended amygdala; VP, ventral pallidum; all other labels as in figures 1 and 3. Tractograms are thresholded with minimum and maximum values equal to 0.5 and 2, respectively.

